# Genome-directed study reveals the diversity of *Salmonella* T6SS effectors and identifies a novel family of lipid-targeting antibacterial toxins

**DOI:** 10.1101/2024.09.27.615498

**Authors:** Gianlucca G. Nicastro, Stephanie Sibinelli-Sousa, Julia Takuno Hespanhol, Thomas W. C. Santos, Joseph P. Munoz, Rosangela S. Santos, Blanca M. Perez-Sepulveda, Sayuri Miyamoto, L. Aravind, Robson F. de Souza, Ethel Bayer-Santos

## Abstract

Bacterial warfare is a common and ancient phenomenon in nature, where bacterial species use strategies to inhibit the growth or kill competitors. This involves the production and deployment of antibacterial toxins that disrupt essential cellular processes in target cells. The continuous arms race in which bacteria acquire new toxin and immunity proteins to promote increased adaptation to their environment is responsible for the diversification of this toxin repertoire. Here, we deployed *in-silico* strategies to analyze 10,000 genomes and identify effectors secreted via the type VI secretion system of *Salmonella*. We identified 128 candidates, which are widespread in a vast array of *Salmonella* serovars and other bacterial species. Tox-Act1 is among the most frequent candidates and was selected for in-depth characterization. Tox-Act1 contains a permuted NlpC/P60 papain-like catalytic core characteristic of lipid-targeting members rather than the typical peptidases or amidases. Evolutionary analysis revealed the relationship of Tox-Act1 with acyltransferases. Biochemical assays with purified toxin and lipidomics of intoxicated cells showed that Tox-Act1 exhibits phospholipase activity, cleaving off acyl groups from phosphatidylglycerol and phosphatidylethanolamine. In addition, we demonstrate that Tox-Act1 is secreted in a T6SS-dependent manner and provide a competitive advantage during colonization of the gut of infected mice. This work broadens our understanding of toxin domains and provides the first direct characterization of a lipid-targeting NlpC/P60 domain in biological conflicts.

## Introduction

Competition is a fundamental biological process in nature, occurring both within and between species that share a common environment. Bacteria actively participate in these ecological battles employing a potent arsenal of toxins as their weaponry[1]. A prominent mechanism for toxin delivery among Gram-negative bacteria is via the type 6 secretion system (T6SS)[2]. Phylogenetic analysis of T6SS components showed that there are four types of T6SSs (T6SS^i-iv^)[3–5], with the canonical T6SS^i^ present in *Proteobacteria* being further classified into six subtypes (i1, i2, i3, i4a, i4b and i5)[4, 6, 7]. The T6SS functions in a contact-dependent manner and relies on the biochemical properties of secreted effector toxins for its function[8]. These toxin domains frequently recombine and move via lateral gene transfer, allowing them to be delivered by different secretion systems, which warrants their name as polymorphic toxins[9].

During secretion via the T6SS, toxins are loaded onto a spear-like structure formed by hexameric rings of Hcp proteins capped by a spike comprising a trimer of VgrG sharpened by a PAAR protein[10]. Effectors are translocated into target cells either fused at the C-terminus of Hcp, VgrG, and PAAR proteins (named specialized effectors), or associated with these proteins via adaptors (cargo effectors)[11]. Antibacterial toxins are often paired with immunity proteins that prevent self-intoxication, thus forming gene pairs that are frequently located near structural components of the T6SS[8, 12].

The protective role of the endogenous microbiota against *Salmonella* infection has been recognized for years[13]; however, only recently have studies started to reveal the mechanism by which the microbiota maintains gut homeostasis and promotes colonization resistance[14]. Despite this understanding, there is a relative scarcity of information about the direct antimicrobial mechanisms employed by commensals and pathogens during these disputes for niche control[15]. T6SSs clusters are conserved across several *Salmonella* spp., highlighting their importance to the fitness of these bacteria[16, 17]. *Salmonella* spp. encode T6SSs in different pathogenicity islands (SPIs), which were acquired by distinct horizontal transfer events[16, 17]. *S.* Typhimurium encodes a T6SS subtype i3 within SPI-6, which is important for interbacterial competition and gut colonization in mice[18, 19]. However, there is limited information about the identity or mechanism of the T6SS effector repertoire that contributes to these phenotypes.

Here, we set out to identify the repertoire of T6SS effectors in a dataset of 10,000 *Salmonella* isolates utilizing a computational approach. Employing sensitive sequence and structure searches alongside comparative genomics, we identified 128 candidates that are widespread among several *Salmonella* serovars and additional bacterial species. This comprehensive analysis indicates that T6SSs subtypes i3 and i1 are associated with antibacterial and anti-eukaryotic effectors, respectively. Furthermore, our findings reveal that each of the 149 serovars harbors a unique combination of effectors, likely required for their specific ecological interactions. A detailed examination of a selected candidate (Tox-Act1) encoding a newly identified domain showed that it is an antibacterial effector belonging to the NlpC/P60 superfamily with a permuted catalytic core. Tox-Act1 is evolutionarily related to acyltransferases and displays phospholipase activity to modify target-cell membrane composition. This study offers a comprehensive characterization of new toxins, especially the arsenal linked to T6SSs in *Salmonella*, identifying novel effector families and providing an in-depth analysis of a new protein family that specifically targets lipids.

## Results

### Computational pipeline for the automatic identification of T6SS components

T6SS effectors are often encoded in close genomic proximity to structural components of the system [16, 20, 21]. To identify new effectors, we employed a "guilt-by-association" approach, which relies on the conservation of genomic context [22]. We applied this methodology to a curated dataset of 10,000 *Salmonella* genomes (10KSG) [23] (Fig 1A). First, we located and classified the T6SS clusters using hidden Markov models (HMMs) from different sources[9, 24–26]. We collected 10 genes upstream and downstream of each locus encoding T6SS components, referred to as genomic sites (Fig 1A). A total of 42,560 genomic sites housing at least one T6SS component were identified. We then applied a graph theoretic strategy leveraging the Jaccard index for network construction[27], followed by the Louvain algorithm[28] to automatically categorize genomic sites. Analysis resulted in five categories: T6SS subtype i1, i2, i3, i4b, and orphan (Fig 1B, 1C).

**Fig 1.**
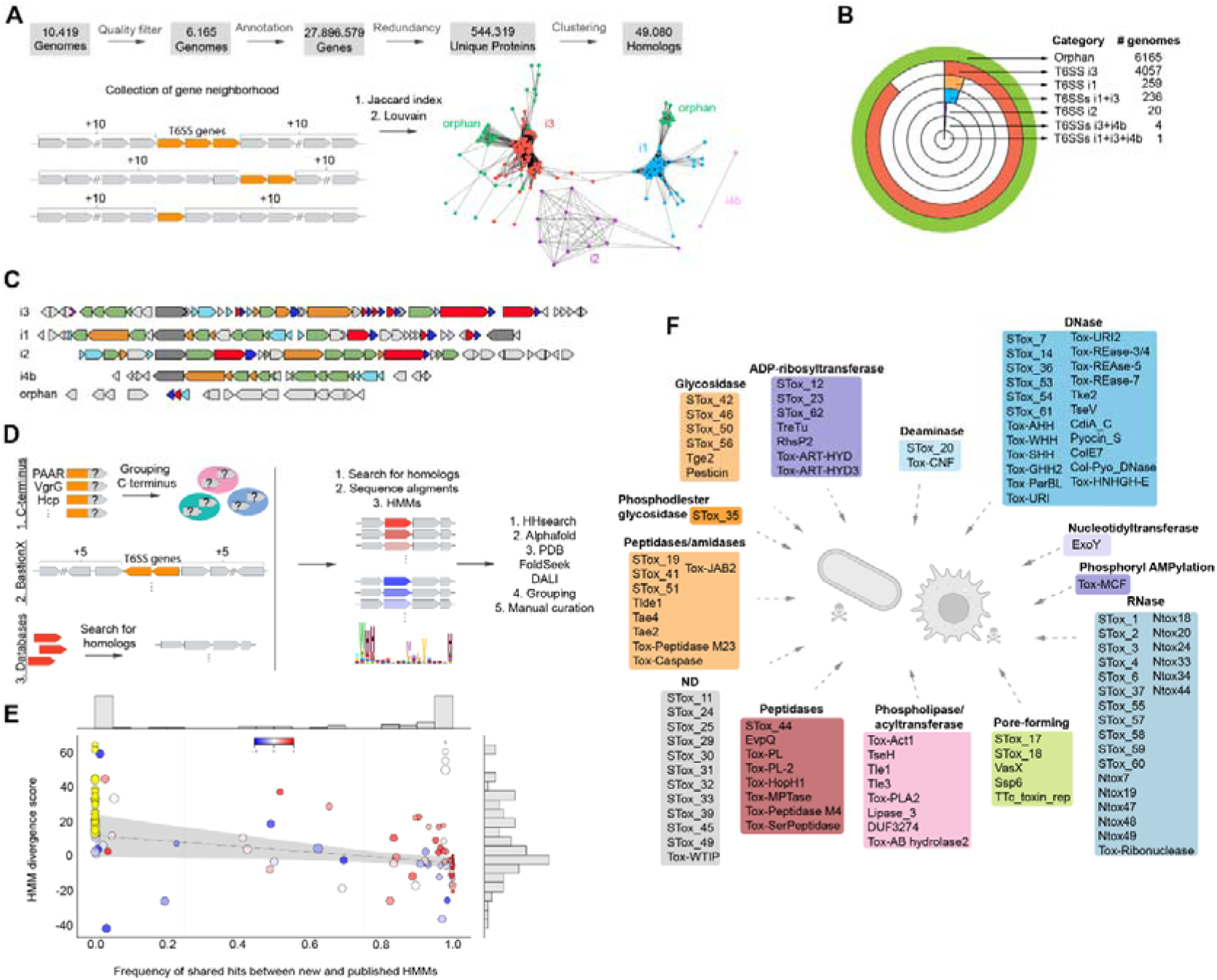
Computational pipeline for the identification of T6SS subtypes and effector repertoire within 10,000 *Salmonella* genomes. **(a)** Pipeline used for classification of genomic sites and T6SS subtypes. **(b)** Number of genomes containing the different T6SS subtypes within the 10KSG dataset. **(c)** Examples of the genomic organization of T6SS structural clusters from distinct phylogenetic subtypes. Colors denote structural proteins forming the membrane complex (orange), sheath and inner tube (light blue), baseplate and spike components (green). Effectors are shown in red and immunities in dark blue. (d) *In silico* strategies used for the identification and classification of T6SS effectors. **(e)** Comparison of profile Hidden Markov Models (pHMMs) of STox to published models of related families. Each circle corresponds to a population of proteins detected by an STox model. The relative frequency of proteins detected by an STox model compared to a reference model is shown on the horizontal axis. The HMM divergence score is shown in the vertical axis. The blue-white-red color scale of each dot represents the values for the Spearman correlation index between STox and reference alignment scores for the same proteins. Yellow circles represent STox models that detect proteins that are not recognized by previously existing models. The radius of each circle is proportional to the total number of proteins detected only by STox models. The regression line does not include the data points for the models represented by yellow circles. **(f)** Schematic representation of the functional classes of T6SS effectors identified in the 10KSG.

We observed mobility of the distinct T6SS subtypes among genomic sites and pathogenicity islands, depending on the strain and/or serovar. Using tRNAs as markers to identify the insertion sites[29], we found that subtype i3 (90%, 4929/5461 genomic sites) is mainly located within SPI-6 and flanked by the aspartate tRNA (tRNA^Asp^)[16], while subtype i1 is not well conserved and found within SPI-19 associated with tRNA^Ala^ (12%, 66/532 genomic sites)[16], and SPI-6 (3%, 18/532 genomes) flanked by tRNA^Asp^ - most subtype i1 could not be assigned to a specific location (82%, 437/532 genomic sites). These results indicate that distinct T6SSs subtypes are inserted into different genomic sites that are hot spots for horizontal transfer events, and the combination between the insertion site and the introduced subtype varies according to the *Salmonella* strain/serovar.

### Identification of the T6SS effectors repertoire in 10,000

#### Salmonella genomes

Next, we focused on identifying effector toxins using three strategies (Fig 1D). First, we focused on proteins containing N-terminal PAAR, VgrG or Hcp domains and additional C-terminal domains with more than 50 amino acids. These C-terminal regions were isolated and grouped based on similarity (80% coverage and 1e-3 e-value). Second, we used the genomic sites containing T6SS components and analyzed up to five genes upstream and downstream via the software BastionX[30]. Third, we used amino acid sequences, HMMs and PSSMs (position-specific scoring matrix) from SecReT6[31], Pfam[25], and Zhang et al. [9] to search the 10KSG dataset for previously described effectors. For candidates not recognized by previously described models, we collected homologs from NCBI using 4 iterations of PSI-BLAST[32] and generated multiple sequence alignments and HMMs.

We then used a series of sequence and structure-based strategies to classify and annotate the function of these candidates: i) profile-profile comparison methods such as HHsearch[33] were used to detect distant homologs; ii) structural models were created using Alphafold2[34] from multiple sequence alignments to perform searches in FoldSeek[35] and DALI[36] against PDB[37] and AlphaFoldDB[38]; iii) the structure-structure comparison algorithm from FoldSeek was used to cluster groups of candidates. All information collected was manually examined to establish the final domain annotation (Fig 1D). Candidates that displayed sequence or structural similarity to known toxin domains or proteins of unknown function that displayed conserved genomic organization and/or adjacent conserved putative immunity proteins across several species were maintained.

In total, we identified 128 groups of effector toxins (Table S1 and Fig 1F). Eighty-thee were already described in public databases (e.g. Ntox47)[9, 25], or represent individual effectors that were experimentally characterized but for which HMMs have not been produced and made available (e.g. TreTu)[21, 39–54]. For the latter, the newly created HMMs were named with a “.st” suffix (e.g. TreTu.st). Within the 128 candidates, 45 groups comprise new toxin domains or highly divergent variations that were not detected by previously published HMMs (Fig 1E). This justified them being distinct groups requiring the design of new HMMs. These groups of effectors were named STox followed by a number (e.g. STox_1) (Table S1 and Fig 1F). It is worth noting that these effectors are not just present in *Salmonella* and are detected widespread across several species, comprising polymorphic toxin domains[9]. A few examples of species encoding each STox, together with the amino acid sequence alignments and AlphaFold prediction can be found at https://leepbioinfo.github.io/10ksgt6ss.

Inspection of genomic context across several bacterial species revealed that most candidates exhibited a conserved adjacent gene coding for a predicted immunity protein, thus suggesting antibacterial activity (83.6%, 107/128). Some effectors, which lacked conserved immunity proteins, were predicted to display anti-eukaryotic activity (12.4%, 14/128) (Table S1 and Fig 1F). The analysis revealed a diverse array of cellular targets and biochemical activities among the 128 toxin groups (Table S1 and Fig 1F). Notably, the activities of a few STox effectors could not be predicted confidently and will require further analysis (Fig 1F). Overall, these findings highlight the significant diversity of effector toxins encoded by *S. enterica* serovars and reveal an array of novel proteins used in interbacterial competition and as virulence factors.

### *Salmonella* serovars encode unique subsets of effector toxins

The most frequent effectors detected within the 10KSG dataset were the peptidoglycan-targeting effectors Tlde1, a L,D-transpeptidase[20]; and Tae4, a papain-like amidase[39] (Fig 2A). These effector families were followed by Ntox47, a predicted RNase with the BECR fold[55]; the metallopeptidase Tox-HopH1; a second more divergent group of Ntox47 (Ntox47.st2); the ADP-ribosyltransferases TreTu[40] and STox_62; and another peptidoglycan-targeting effector Tae2 [39] (Fig 2A). Together, these eight effectors were found in most genomes within the 10KSG and constitute the core T6SS effectors. In addition, by analyzing the combination of effectors in each genome, we found that 88% of the genomes within the 10KSG dataset were estimated to encode up to 4 effectors per genome (mode = 3), while 12% encode combinations that range between 5-18 effectors (mode = 5) (Fig S1).

**Fig 2.**
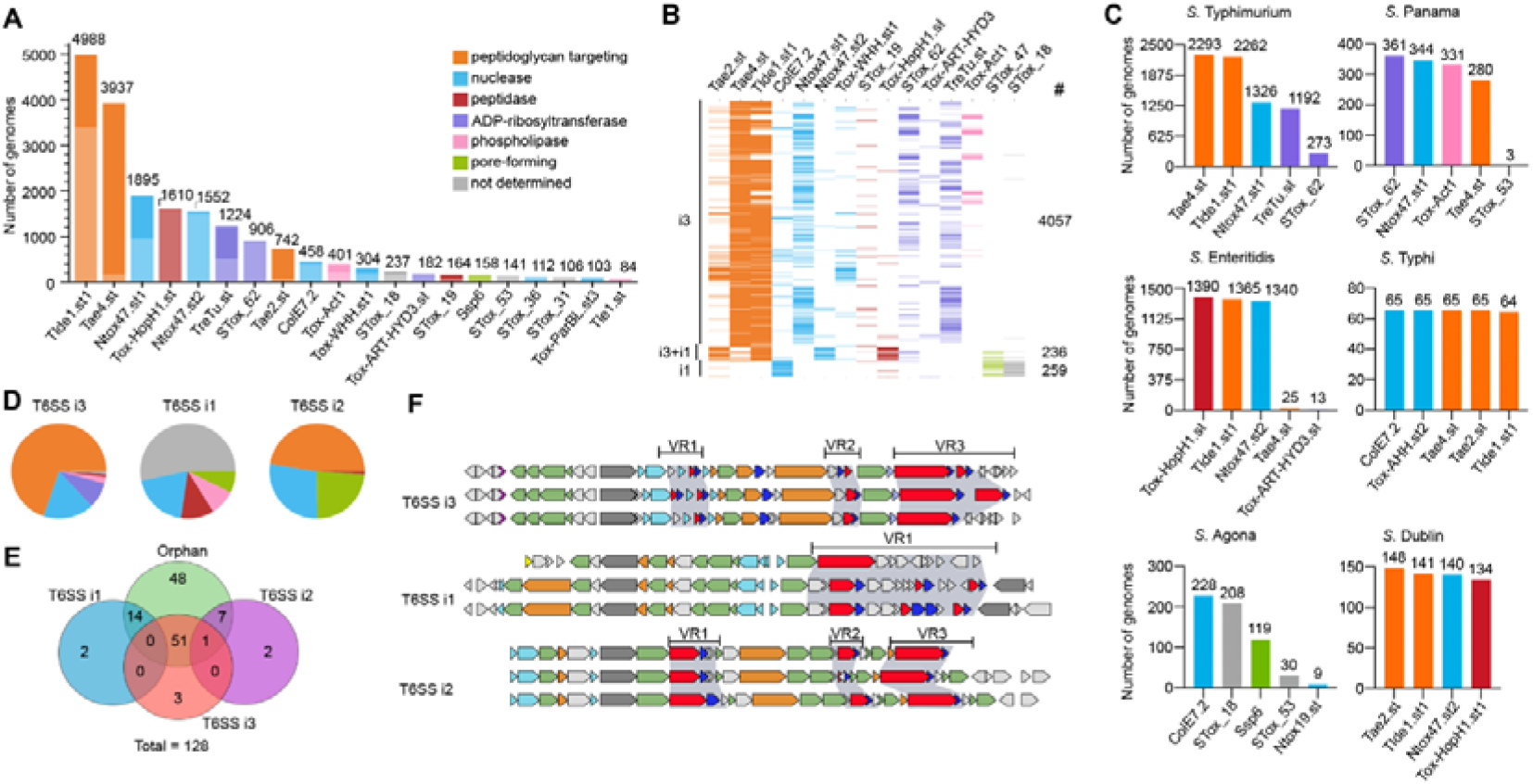
Unique subsets of effectors are associated with specific *Salmonella* serovars and T6SS subtypes. **(a)** The most frequent effectors identified in the 10KSG dataset. Each bar represents the number of genomes encoding a specific effector. Colors represent different effector activities, with light colors representing orphan effectors while dark colors represent effectors encoded within the structural cluster. **(b)** Schematic representation of the most common sets of effectors in genomes encoding different T6SS subtypes. The number of genomes is indicated on the right. Colors represent activity as shown in (a). **(c)** The five most frequent effectors encoded in different *Salmonella* serovars. Colors indicate the effector activity as in (a). **(d)** Pie chart illustrating the relative proportions of effectors classified by activity encoded within the T6SSs subtypes i3, i1, and i2. Colors as shown in (a). **(e)** Venn diagram illustrating the proportion of overlap between effectors encoded within each T6SS structural cluster (blue: i1; purple: subtype i2; red: subtype i3; and green: orphan). **(f)** Schematic representation of the genetic organization of T6SSs showing the position of variable regions in which the effector and immunity proteins are encoded. Colors denote structural proteins forming the membrane complex (orange), sheath and inner tube (light blue), baseplate and spike components (green). Effectors are shown in red, and immunities in dark blue.

Next, we determined the 5 most frequent effectors detected in each of the 149 *Salmonella* serovars. Serovars that predominantly encode subtype i3, such as *S.* Typhimurium. *S.* Panama, *S.* Infantis, and *S.* Typhi, harbor effectors targeting peptidoglycan (e.g. Tae4, Tlde1, Tae2), nucleases (e.g. Ntox47, Tox-WHH, ColE7, Tox-AHH) and ART superfamily enzymes (e.g. TreTu, STox_62) (Fig 2B, 2C). *S.* Dublin, which contains both subtypes i1 and i3, displays a mixture of effectors targeting the peptidoglycan (Tae2 and Tlde1), a nuclease (Ntox47), and a metallopeptidase (Tox-HopH1) (Fig 2C). *S.* Agona encodes only a T6SS subtype i1 and contains effectors with nuclease (ColE7 and Ntox19), pore-forming (STox_47), and undetermined activities (STox_18 and STox_53) (Fig 2C). The core effectors of each serovar can be found in Fig S2.

### Each *Salmonella* T6SS subtype is associated with target-specific effectors

We analyzed the combination of effectors most frequently encoded in genomes containing phylogenetically distinct T6SSs. Our findings revealed that the antibacterial subtype i3 predominantly displays a combination of peptidoglycan-targeting effectors (e.g. Tae4 and Tlde1); nucleases (e.g. Ntox47 and Tox-WHH); and ART family enzymes (e.g. TreTu and STox_62) (Fig 2A, 2B). Conversely, genomes encoding the anti-eukaryotic subtype i1 lack peptidoglycan-targeting effectors and show a combination of nucleases (e.g. ColE7), pore-forming toxins (e.g. Ssp6) and effectors with undetermined activity (Fig 2B, 2C). For genomes encoding both subtypes i1 and i3, there was a mix of effectors with antibacterial (e.g. Tae2, Tlde1, Ntox47) and anti-eukaryotic activity (e.g. Tox-HopH1) (Fig 2B, 2C). This data supports the classification of *Salmonella* T6SS subtype i3 as antibacterial and subtype i1 as anti-eukaryotic.

Notably, effectors encoded within or close to T6SS structural clusters, show minimum overlap (Fig 2E), suggesting an evolutionary scenario where T6SS clusters are acquired alongside with the associated set of effectors and/or that certain subsets of effectors are preferably exchanged among bacteria harboring similar T6SS subtypes. Previous analysis of subtype i3 identified the insertion of thee variable regions between the structural genes (VR1-3) in which effectors are encoded [16] (Fig 2F). Our results indicate that VR1 and VR2 contain mainly toxins targeting the periplasm (e.g. Tae2, Tae4, Tlde1, Tox-Act1) whereas VR3 primarily houses toxins targeting the cytoplasm (e.g. Ntox47, Tox-WHH, TreTu). The latter are typically associated with an N-terminal PAAR domain and Rhs (Rearrangement hotspot) repeats, which assist in the translocation across the bacterial inner membrane to reach the cytoplasmic targets[40, 55–57] (Fig 2F). The position of effectors at the edge of the T6SS cluster and the domain architecture containing Rhs repeats facilitate recombination events[55], possibly leading to the greater diversity of nuclease effectors observed at the VR3 (Fig S1).

### Tox-Act1 is a T6SS effector used for interbacterial competition in the mouse gut

Among the newly identified effectors, STox_15, which was renamed Tox-Act1 (toxin acyltransferase 1, see details below), emerged as one of the most abundant (Fig 2A). To analyze whether Tox-Act1 and its associated downstream gene (Imm-Act1) form an effector and immunity pair (Fig 3A), we cloned these genes in compatible vectors under the control of different promoters and assessed toxicity upon expression in *Escherichia coli*. Tox-Act1 is toxic in the periplasm (SP-Tox-Act1) but not in the cytoplasm (Tox-Act1) of *E. coli* and co-expression with Imm-Act1 neutralizes the effect (Fig 3B).

**Fig 3.**
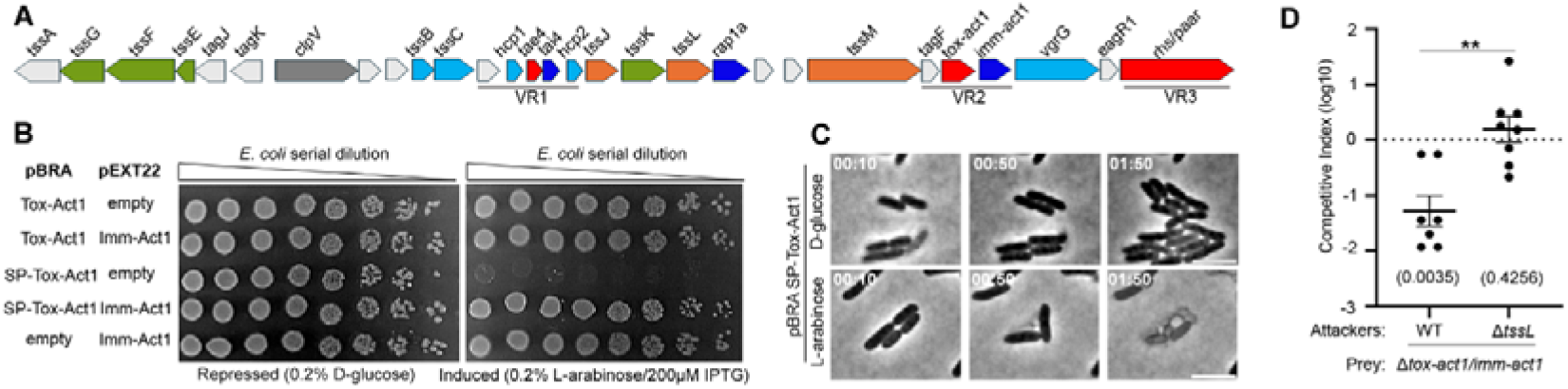
Tox-Act1 is a T6SS effector used for bacterial competition in the mouse gut. **(a)** Scheme of the genomic region encoding Tox-Act1 and Imm-Act1 effector/immunity pair (FD01843896). **(b)** *E. coli* toxicity assay. Serial dilutions of *E. coli* carrying pBRA and pEXT22 constructs. Images are representative of thee independent experiments. **(c)** Time-lapse microscopy of *E. coli* carrying pBRA SP-Tox-Act1 grown on repressed or induced conditions. Scale bar: 5 µm. Timestamps in hh:mm. **(d)** C57BL/6 mice were infected by oral gavage with equal numbers of each strain. Bacteria were recovered from the cecum 4 days after infection, and CI values calculated. The log10 Cis were used for statistical analysis. Single sample t-test was used to compare the CI to the hypothetical value of 0, and *p* value is indicated in brackets. Unpaired t-test (\*\**p* <0.01) was used to compare the two groups.

We performed time-lapse microscopy to evaluate bacterial growth and morphology at the single cell level. *E. coli* carrying the plasmid with SP-Tox-Act1 grew normally in d-glucose (repressed conditions) (Movie S1); however, shortly after induction of SP-Tox-Act1 with l-arabinose, cells began lysing (Movie S2). It was curious that cells lysed without losing their rod shape, which suggests that the peptidoglycan was not affected (Fig 3C). After lysing, residual structures resembling the intact peptidoglycan sacculus remained (Fig 3C), indicating that this is not the target of Tox-Act1. In addition, we noticed that the cognate immunity protein Imm-Act1 contains a conserved domain with two transmembrane helices (Fig S3), suggesting that the site of its neutralizing action occurs at the cell membrane.

The SPI-6 T6SS of *Salmonella enterica* serovar Typhimurium is repressed by the silencer protein H-NS [18]. Bacterial competition assays performed *in vitro* have not been able to recapitulate the natural conditions found in the mouse gut to fully activate this system [19]. To analyze whether Tox-Act1 is a T6SS substrate and contributes to interbacterial competition *in vivo*, we performed competitive index assays using *S.* Typhimurium strain IC33Q that naturally encodes Tox-Act1 (Fig 3A). Mice were infected by oral gavage with a 1:1 mixture of wild-type (WT) and the effector-immunity double mutant Δ*tox-act1/imm-act1*, or a mixture of the T6SS mutant (Δ*tssL*) and Δ*tox-act1/imm-act1*. Results showed that Δ*tox-act1/imm-act1* has a competitive disadvantage during colonization of the mouse gut when compared to the WT, and such disadvantage is not observed when compared to Δ*tssL* (Fig 3D). These results confirm that Tox-Act1 is an antibacterial effector secreted via the SPI-6 T6SS and that it is actively expressed by this *S. enterica* strain and used during competition in the mouse gut.

### Tox-Act1 is evolutionarily related to NlpC/P60 enzymes targeting lipids

The Tox-Act1 domain was not identified by any of the HMMs deployed in the initial steps of this study. However, subsequent HHpred analysis revealed a significant probability of homology with DUF4105 and the effector TseH from *Vibrio cholerae*[45] (data not shown), both of which are members of the NlpC/P60 superfamily[58, 59]. This superfamily was previously defined as encompassing four families, which are divided into two higher-order groups (canonical and permuted)[58].

Members of the canonical group (AcmB-like and P60-like) function as peptidases involved in peptidoglycan hydrolysis[60], while permuted members (YaeF-like and LRAT-like) exhibit a circular permutation in their catalytic core, creating a hydrophobic binding pocket that provides specificity for lipids [61, 62]. Closer inspection of the sequence and structure of Tox-Act1 revealed a circular permutation of the catalytic domain indicating that it belongs to the permuted NlpC/P60 group (https://leepbioinfo.github.io/10ksgt6ss/)[58].

To predict the function of Tox-Act1, we sought to understand its evolutionary relationship by constructing a phylogenetic tree using the sequences of Tox-Act1, TseH and additional permuted members, such as LRAT and YiiX. The analysis revealed the formation of five major clades including YiiX, LRAT, TseH, Tox-Act1 and DUF4105 (Fig 4A, Table S2 and Table S3). Genomic context analysis predicted that proteins of the Tox-Act1 clade are toxins deployed in biological conflicts (Fig 4A and 4B). Similarly, the TseH clade (Pfam DUF6695) displays genomic contexts indicative of biological conflicts (Fig 4A and 4B). The clades LRAT and YiiX harbor proteins known to be involved in lipid metabolism: LRAT (lecithin:retinol acyltransferase) is an enzyme present in mammals and involved in the transference of acyl groups from phosphatidylcholine to all-trans retinol to produce all-trans retinyl esters that are storage forms of vitamin A[63]; H-RAS-like suppressor (HASLS) proteins are a group within the LRAT family that display both acyltransferase and phospholipase A1/2 activities[64]; and YiiX-like family members from *Bacillus cereus* are active against lipids[62]. We decided to name Tox-Act1 (toxin acyltransferase 1) due to its evolutionary relationship with acyltransferases.

**Fig 4.**
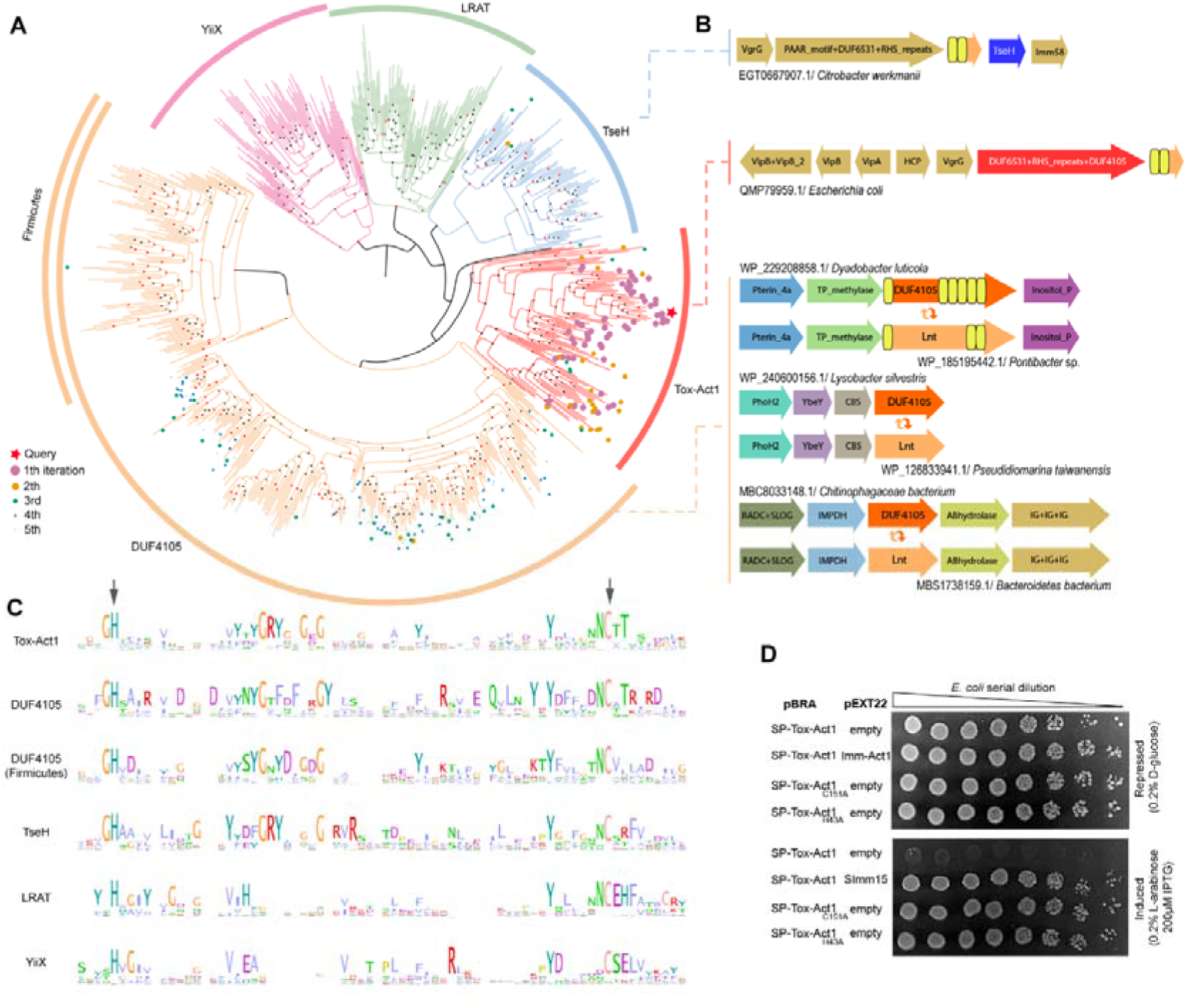
Tox-Act1 is evolutionarily related to lipid-targeting enzymes with a permuted NlpC/P60 domain. **(a)** Maximum-likelihood phylogenetic tree of permuted NlpC/P60 members. Dots represent the number of PSI-BLAST iterations required to collect homologs and the red star marks the query. **(b)** Genomic organization of representatives from clades TseH and Tox-Act1 showing the genes are encoded in the context of conflict systems, and DUF4105 showing context of lipid metabolism. **(c)** Sequence logo from the enzymatic core of permuted NlpC/P60 from all clades shown in (a). The arrows indicate conserved His and Cys residues that were mutated in (d). **(d)** *E. coli* toxicity assay. Serial dilution of *E. coli* containing pBRA and pEXT22 constructs. Images are representative of thee independent experiments.

Interestingly, Tox-Act1 emerged as the sister clade of DUF4105 (Fig 4A and 4B). Our comparative genomics analysis revealed a recurring evolutionary pattern in which DUF4105 domain-containing proteins are repeatedly displaced by apolipoprotein N-acyltransferases (Lnt) across thee distinct genomic contexts (Fig 4B). Hence, we proposed that DUF4105 could be working as an acyltransferase. Remarkably, DUF4105 was recently identified as the missing lipoprotein N-acyltransferase in *Bacteroides*[65], which was named Lnb (N-acyltransferase in *Bacteroides*). These experimental results confirmed our independent *in-silico* prediction for the function of DUF4105 and provide two different lines of evidence leading to similar conclusions. It is noteworthy that the DUF4105 clade identified in our analysis consists primarily, though not exclusively, of *Bacteroides* species, with a branch enriched in Gram-positives like *Firmicutes* (Fig 3A). The list of homologs containing DUF4105 can be found in Table S3C and S3D.

Multiple sequence alignments of each of the permuted clades including Tox-Act1 revealed the conserved catalytic His and Cys residues characteristic of the NlpC/P60 superfamily (Fig 4C)[58]. Substitution of these residues for alanine (Tox-Act1_H43A_ and Tox-Act1_C151A_) eliminated toxicity in *E. coli* (Fig 4D). These findings collectively confirm that the enzymatic function of the NlpC/P60 papain-like fold domain is crucial for toxicity. Collectively, the periplasmic-acting phenotype of Tox-Act1, the presence of a membrane-associated immunity protein and the fatty acyl linkage targeting activities common in the permuted NlpC/P60 members strongly support a function for Tox-Act1 in targeting membrane lipids.

### Tox-Act1 displays phospholipase activity and changes the membrane composition of intoxicated cells

To analyze the enzymatic activity of Tox-Act1, we incubated purified recombinant protein (Fig S4) with either purified phosphatidylglycerol (PG) 16:0-18:1 or phosphatidylethanolamine (PE) 16:0-18:1 and analyzed the reaction product by HPLC coupled to mass spectrometry (Fig 5A and B). Results showed that Tox-Act1 has predominantly phospholipase A1 activity and cleaves both PG and PE (a preference for cleaving off the 16:0 acyl chain) as observed by the accumulation of 18:1 lysophospholipids (Fig 5A and B).

**Fig 5.**
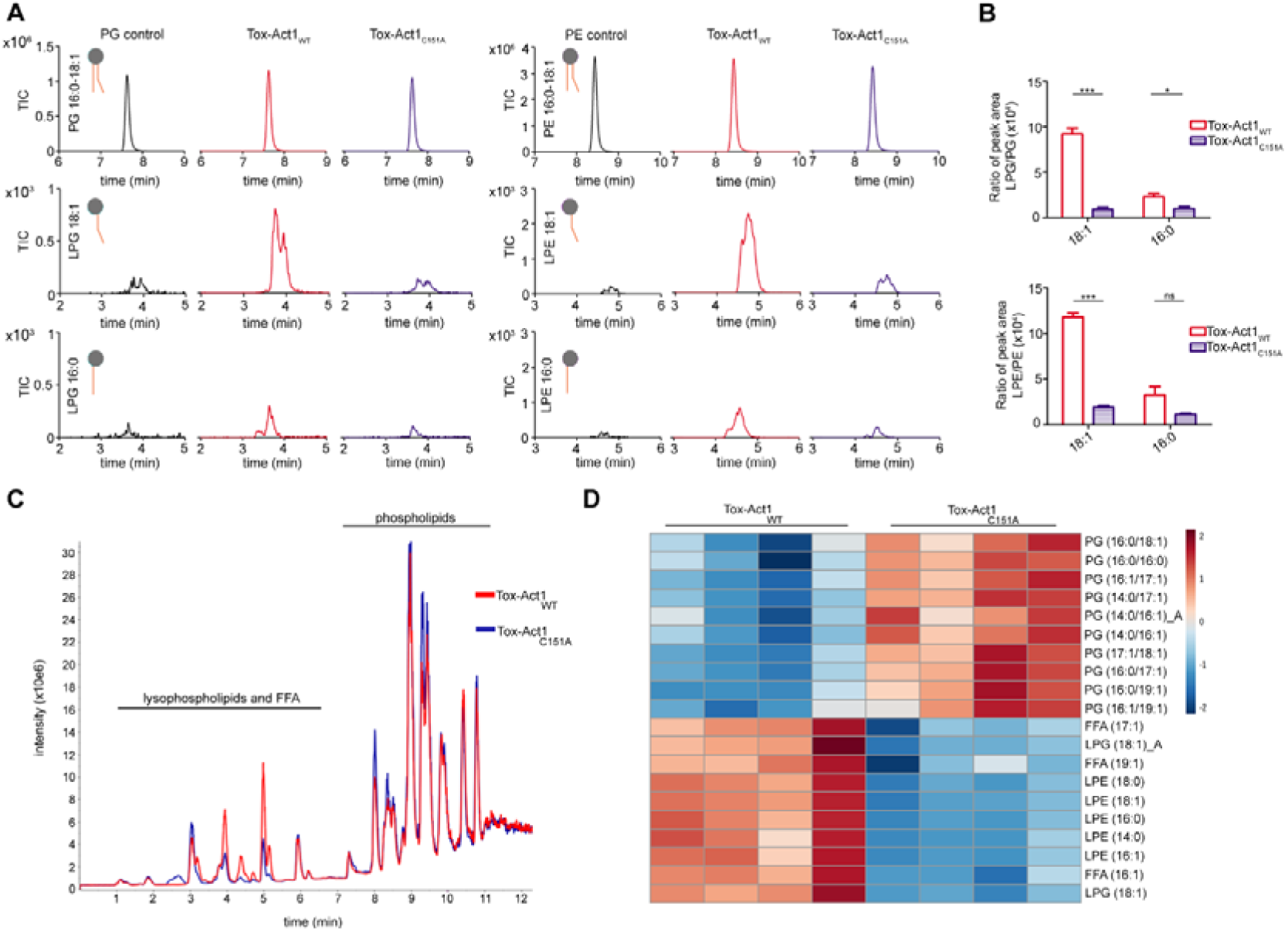
Tox-Act1 has phospholipase A activity and changes the composition of target cell membranes. **(a)** *In vitro* enzymatic assay with recombinant Tox-Act1 (red) or Tox-Act1_C151A_ (blue) incubated with different phospholipids (both 16:0-18:1 PG and PE). The amount of lysophospholipids produced was analyzed and quantified by HPLC-MS/MS. **(b)** Quantification of the peak area of lysophospholipids normalized by the intact substrate. Data corresponds to the mean ± SD. ***p < 0.001 and *p < 0.05, *ns* not significant (unpaired t-test). **(c)** UHPLC-MS total ion chomatogram showing the profile of total lipids extracted from *E. coli* expressing Tox-Act1 (red) or Tox-Act1_C151A_ (blue). **(d)** Heatmap plot of top 20 altered lipids of intoxicated *E. coli*. Results display four biological replicates of each condition (WT or C151A) with the quantification of lipids species (Tukey test; p < 0.05 FDR adjusted). Data were expressed in nM mg of protein^-1^ and normalized by log transformation (base 10) prior to analysis. Red (up) and blue (down) bars represent changes in lipid species concentration relative to the normalized mean. Letters differentiate between the isomers.

Next, we set out to determine whether Tox-Act1 could cause changes in the composition of phospholipids when ectopically expressed by target cells. *E. coli* harboring the SP-Tox-Act1 plasmid were grown to OD_600nm_ of 1.0 in the presence of d-glucose (repressed), washed and resuspended in AB medium with l-arabinose to induce the expression of the toxin. Total lipids were extracted and analyzed by UHPLC-MS. We observed a general decrease in intact phospholipid forms in Tox-Act1_WT_, especially phosphatidylglycerol (PG), when compared with the catalytic mutant Tox-Act1_C151A_ (Fig 5A and 5B). In addition, an increase in lysophospholipids forms - either lysophosphatidylglycerol (LPG) or lysophosphatidylethanolamine (LPE) - and free fatty acids (FFA) was detected in the wild-type (Fig 5A and 5B). Lysophospholipids possess amphiphilic properties and have an inverted cone-shaped molecular structure that interacts with and modifies membrane properties (e.g. curvature) similarly to detergents[66]. The accumulation of lysophospholipids on the target cell membrane likely promotes the observed membrane disruption (Fig 3C) due to its detergent-like properties[67]. Collectively, these results confirm that Tox-Act1 targets phospholipids similarly to other proteins possessing a permuted NlpC/P60 domain, and that Tox-Act1 displays phospholipid-degrading activity. It remains to be determined whether Tox- Act1 also possesses acyltransferase activity as reported for other permuted NlpC/P60.

## Discussion

Our comprehensive analysis of 10,000 *Salmonella* genomes has significantly expanded the repertoire of T6SS effectors and polymorphic toxins in general, identifying 128 candidates, among which 45 comprise novel protein domains. Our study employed a robust bioinformatic pipeline, integrating classical methods with the latest structural bioinformatics techniques. The combination of sensitive sequence and structure searches with comparative genomics provided a comprehensive understanding of the identified toxin domains. In addition, the manual curation of candidates ensured high confidence in our results, distinguishing our approach from previous large-scale *in-silico* analyses. The identification of these novel toxin domains builds upon the foundational work by Zhang et al. [9] and others [30, 68–76], which characterized the diversity of polymorphic toxin systems across bacterial lineages. The identification of novel toxin domains highlights the constant evolutionary arms race between bacteria, driving the diversification of toxin and immunity proteins.

This study not only broadens our understanding of *Salmonella* T6SS effectors and toxin domains but also provides the first direct characterization of a lipid-targeting NlpC/P60 domain. The phospholipase activity of Tox-Act1, which cleaves acyl groups from phosphatidylglycerol and phosphatidylethanolamine, underscores its role in membrane disruption during bacterial competition. Interestingly, our phylogenetic analysis revealed an evolutionary link between Tox-Act1 and a new family of lipoprotein N-acyltransferases in *Bacteroides*, adding a new dimension to the functional diversity of these toxins. Notably, one of the homologs of Tox-Act1 is TseH [45, 77], which has been proposed to be an endopeptidase due to its NlpC/P60 domain and similarity to the amidase Tse1[78, 79]. However, unlike Tse1, TseH exhibits a permutation in its catalytic core [77]. This permutation, along with its evolutionary relationship to Tox-Act1 and other permuted NlpC/P60, suggest that TseH actual substrate might be an acyl group in phospholipids rather than a peptide/amide bond. Exploring the potential acyltransferase activity of Tox-Act1 and its homologs could reveal further biochemical diversity within the NlpC/P60 superfamily.

In the context of *Salmonella* biology, the unprecedented diversity of T6SS effectors presents numerous opportunities for new studies. Our findings reveal the existence of individual subsets of T6SS effectors for each serovar, suggesting that *Salmonella* acquire and maintain effectors in response to specific environmental pressures rather than accumulating an increasingly larger array of effectors. Notably, we observed a higher number of effectors in serovars isolated from environmental sources compared to those from patients, indicating that the number of effectors increases in more diverse environments where there are potentially more encounters with a variety of rival species. In addition, we observed that *Salmonella* serovars encode phylogenetically distinct T6SS clusters, which are specialized to target either eukaryotic or bacterial cells. These results further highlight the versatility of the T6SS aiding in adaptation to many environments and hosts.

In conclusion, our comprehensive analysis has greatly enhanced the understanding of toxins involved in bacterial competition and pathogenesis. The identification of previously uncharacterized toxin domains highlights the potential for discovering novel biochemical activities. This study provides a solid foundation for future research into the complex dynamics of conflict systems and their implications for bacterial ecology and pathogenesis.

## Materials and Methods

### Comparative genomic analysis

The .gff files from the genome assemblies retrieved from the 10KSG project[23] were organized and stored in tabular format using Python scripts, based on the Biopython[80] and pandas[81] libraries. Iterative searches were conducted using Jackhmmer[82] with a 10e-6 e-value cutoff. Protein clustering was performed using MMseqs[83] to remove redundancy (80% coverage and 70% identity) and form homologous groups (80% coverage and e-value 10e-3). Multiple sequence alignments were generated using the local-pair algorithm in MAFFT[84], and phylogenetic trees were constructed using FastTree[85]. Domain identification and annotation was performed using HMMsearch and HMMscan[82, 86] and models from the databases Pfam[25], TXSScan[24], and BastionHub[87]. Remote homology identification was performed using HHpred[33] and FoldSeek.

### Bacterial strains

A list of bacterial strains used in this work can be found in Table S4. Strains were grown at 37 °C in Lysogeny Broth (10 g/L tryptone, 10 g/L NaCl, 5 g/L yeast extract) under agitation. AB medium was used for lipidomics: 0.2% (NH4)2SO2, 0.6% Na2HPO4, 0.3% KH2PO4, 0.3% NaCl, 0.1 mM CaCL2, 1 mM MgCl2, 3 μM FeCL3, supplemented with 0.2% sucrose, 0.2% casamino acids, 10 μg/mL thiamine, and 25 μg/mL uracil. Cultures were supplemented with antibiotics in the following concentration when necessary: 50 μg/mL kanamycin, 100 μg/mL ampicillin, and 50 μg/mL streptomycin.

### Cloning and mutagenesis

All primers are listed in Table S4. Tox-Act1 and Imm-Act1 were amplified by PCR and cloned into pBRA vector under the control of P_BAD_ promoter[88] with or without pelB signal peptide sequence from pET22b (Novagen)[89]. Imm-Act1 was cloned into pEXT22 under the control of P_TAC_ promoter[90]. For protein expression and purification, Tox-Act1 was cloned into pET28a (Novagen), including a C-terminal Strep II tag. Point mutations (Tox-Act1_H43A_, Tox-Act1_C151A_) were created using QuikChange II XL Site-Directed Mutagenesis Kit (Agilent Technologies) or by splicing by overlap extension (SOE) PCR. All constructs were confirmed by sequencing. *S*. Typhimurium mutant strains used for competition assays were constructed by λ-Red recombination system using a one-step inactivation procedure [91].

### *E. coli* toxicity assay

Overnight cultures of *E. coli* DH5α co-expressing effectors for cytoplasmic (pBRA Tox-Act1) or periplasmic (pBRA SP-Tox-Act1) localization and immunity protein (pEXT22 Imm-Act1) were adjusted to OD_600nm_ 1, serially diluted in LB (1:4) and 5 μL were spotted onto LB agar (1.5%) containing either 0.2% d-glucose or 0.2% l-arabinose plus 200 μM IPTG, supplemented with streptomycin and kanamycin, and incubated at 37°C for 20 h.

### Time-lapse microscopy

For time-lapse microscopy, LB agar (1.5%) pads were prepared by cutting a rectangular piece out of a double-sided adhesive tape, which was taped onto a microscopy slide as described previously[89]. *E. coli* DH5α harboring pBRA SP-Tox-Act1 was subcultured in LB (1:50) with 0.2% d-glucose until reaching OD_600nm_ 0.4–0.6 and adjusted to OD_600nm_ 1. Cultures were spotted onto LB agar pads supplemented either with 0.2% d-glucose or 0.2% l-arabinose plus antibiotics. Images were acquired every 15 min for 16 h using a Leica DMi-8 epifluorescent microscope fitted with a DFC365 FX camera (Leica) and Plan-Apochomat ×63 oil objective (HC PL APO ×63/1.4 Oil ph3 objective Leica). Images were analyzed using FIJI software [92].

### Competitive index

Female C57BL/6 mice (6-8 weeks old) were purchased from Jackson’s Laboratory. Mice were housed in pathogenic free conditions with unlimited access to food and water, except for 4 hours prior oral gavages. Mice were pre-treated with 20 mg of streptomycin 24 hours prior infection with *Salmonella*. Mice were infected by oral gavage with a 1:1 mixture (total of 10^10^ CFUs) of *S.* Typhimurium wild-type (WT) and Δ*tox-act1/imm-act1* (Cm^R^), or a mixture of Δ*tssL* and Δ*tox-act1/imm-act1*. Mice were euthanized four days after infection by CO_2_ asphyxiation, and cecum were harvested and the content serial diluted in PBS 1x (Phosphate-buffered saline) and plated in MacConkey Agar to determine the total CFU counts. One hundred colonies from each mouse were patched on chloramphenicol plates to determine the proportion of each strain in the mixture. The same procedure was performed with the initial mixture prior infection. Competitive index was calculated as previously described [93]. The unpaired t-test was used to compare the CI between WT and Δ*tssL* groups, while the single sample t-test was used to compare each log10 CI to the hypothetical value of 0 (the value of 0 means that two strains grew equally well *in vivo*).

### Protein expression and purification

*E. coli* BL21(DE3) carrying pET28a Tox-Act1_WT_-Strep or Tox-Act1_C151A_-Strep were grown in 4 L of LB supplemented with kanamycin (37 °C, 180 rpm) until OD_600nm_ 0.7. Expression was induced with 1 mM IPTG for 16 h at 16 °C. Cells were harvested via centrifugation at 5000 *g* for 20 min, and pellets were resuspended in lysis buffer (50 mM Tris HCl pH 8.0, 350 mM NaCl, 45 mM β-mercaptoethanol, 5 mg/mL lysozyme, 10% glycerol) and lysed at 4°C using a sonicator. The lysate was centrifuged at 40,000 *g* for 45 min at 4°C. The supernatant was loaded onto a 1 ml StrepTrap HP column (Cytiva) equilibrated in buffer (50 mM Tris-HCl pH 8.0, 350 mM NaCl, 10% glycerol). The column was washed with 40 column volumes (CV) of wash buffer (50 mM Tris-HCl pH 8.0, 1 M NaCl, 10% glycerol, 45 mM β-mercaptoethanol), followed by a second wash with 12 CV (50 mM Tris-HCl pH 8.0, 1.5 M urea) to remove chaperonin GroEL[94]. The column was subjected to a third round of washes with 40 CV of wash buffer and eluted with 10 CV of elution buffer (50 mM Tris-HCl pH 8.0; 350 mM NaCl; 10% glycerol; 50 mM biotin). Fractions were buffer exchanged (25mM Tris-HCl pH 7.5, 100 mM NaCl, 5% glycerol) and concentrated using an Amicon of 30 kDa (Sigma). Protein aliquots were snap-frozen until use.

### *In vitro* phospholipase assay

For *in vitro* enzymatic assay, phospholipids 1-palmitoyl-2-oleoyl-sn-glycero-3-phospho-(1’-rac-glycerol) (PG 16:0-18:1) and 1-palmitoyl-2-oleoyl-sn-glycero-3-phosphoethanolamine (PE 16:0-18:1) were purchased from Avanti Polar Lipids. Substrates were resuspended and diluted in methanol to adjust the concentration of aliquots. The methanol of each aliquot was dried under a nitrogen flow. A total of 1.2 mM of phospholipids (PG or PE) were resuspended in reaction buffer (25 mM Tris-HCl pH 7.5, 100 mM NaCl, 0.5 mM CaCl_2_, 0.5 mM MgCl_2_, 180 mM sodium deoxycholate and 0.5 mM DTT) and incubated with 0.8 mM of enzyme (Tox-Act1_WT_ or Tox-Act1_C151A_) in a total volume of 100 µL for 2 h at 37°C under agitation (350 rpm). Lipids were extracted by adding 830 µL of a mixture of MTBE/methanol/water (10:3:2.5, v/v/v), followed by incubation under agitation for 1 h at room temperature. Samples were centrifuged for 2 min at 220 *g* and 350 µL of the top fraction was transferred to a new tube, dried in a SpeedVac and stored at –80 °C until analysis.

For mass spectrometry analysis, samples were resuspended in 350 µL of isopropanol and analyzed in a Shimadzu 8060 Triple Quadrupole Liquid Chomatograph Mass Spectrometer. Samples (0.1-0.5 µL) were loaded into an Agilent column C18 ZORBAX Eclipse Plus (4.6 x 150 mm, 5 µm, 400 bar) with a flow rate of 0.5 mL/min and an oven temperature of 40 °C. HPLC gradients were as described below for lipidomic analysis. The phospholipids and lysophospholipids of interest were analyzed in the multiple reaction monitoring (MRM) mode using m/z transitions, collision energies (CE) and dwell times as shown in Table S5A. Data was acquired by Shimadzu LabSolutions and processed in LabSolutions Browser. Graphs were plotted using GraphPrism 5.

### Lipidomics of Tox-Act1-intoxicated *E. coli*

*E. coli* MG1655 containing the plasmids pBRA SP-Tox-Act1_WT_ or SP-Tox-Act1_C151A_ were cultured in LB containing 0.2% d-glucose at 37 °C for 14 h. Cells were subcultured in LB 0.2% d-glucose until an OD_600nm_ of 1 and centrifuged (10 min, 2900 *g*, 30 °C). Cells were washed with 40 mL of preheated AB medium at 37 °C, centrifuged (10 min, 2900 *g*, 30 °C), and resuspended in 5 mL of AB supplemented with 0.2% l-arabinose to induce Tox-Act1 expression. Cells were incubated for 1 h at 37 °C with agitation (100 rpm). The cells were centrifuged (15 min, 2900 *g*, 4 °C) and washed once with 1 mL of PBS pH 7.4. PBS was removed by centrifugation and the cell pellet was stored at -80 °C until lipid extraction. A cocktail of class specific internal standards was added to the cell mass prior to lipid extraction for subsequent quantification and normalization (Table S5B). Total lipid extraction was performed using an adapted version of the protocol described by Yoshida, Kodai (95). Briefly, cell pellets were resuspended with 500 µL of cold methanol and 1 mL of MilliQ water and transferred to glass tubes. A mixture of chloroform and ethyl acetate (4:1) was added, followed by agitation for 1 min at 25 °C. Samples were centrifuged (2 min, 1500 *g*, 4°C) and the lipid-containing phase (lower phase) was extracted and transferred to a new glass tube that was dried under a nitrogen (N_2_) flow until all solvent traces were evaporated. Samples were stored at -80 °C until analysis.

Lipid extracts were diluted in 100 µL of isopropanol and analyzed using ultra-high performance liquid chomatography (UHPLC Nexera, Shimadzu) coupled with an ESI-Q-TOF mass spectrometer (Triple TOF 6600, Sciex) (UHPLC-Q-TOF/MS). 2 μL of each sample were injected into the UHPLC-MS, and molecules were separated using a CORTECS column (C18, 1.6 μm, 2.1x100 mm, Waters) with a flow rate of 0.2 mL/min and temperature set to 35 °C [96]. The mobile phases consisted of (A) water/acetonitrile (60:40) and (B) isopropanol/acetonitrile/water (88:10:2). Ammonium acetate at a final concentration of 10 mM was incorporated in both mobile phases A and B for the negative ionization acquisition mode. The gradient elution used in the chomatography was from 40 to 100% (mobile phase) B over the first 10 min; 100% B from 10-12 min; 100 to 40% B for 12-13 and holding 40% B for 13-20 min. The negative mode was utilized for the examination of phospholipids and free fatty acids. MS and MS/MS data acquisition was performed using Analyst 1.7.1 software (Sciex). Mass spectrometry data was inspected using PeakView 2.0 software (Sciex), and lipid molecular species were manually identified with the help of an in-house manufactured Excel-based macro. Lipid species were quantified using MultiQuant software (Sciex), where the precursor ions areas were normalized by the internal standards for each class (Table S5B).

## Data availability

All data supporting the findings of this study are available within the paper and its Supplementary Information files or at https://leepbioinfo.github.io/10ksgt6ss/. All data and code used for sequence and genome context analyses are available on a GitHub repository at https://github.com/leepbioinfo/10ksgt6ss. The ROTIFER package can be downloaded from the GitHub repository https://github.com/leepbioinfo/rotifer. The contents of both repositories are made available under the Creative Commons Attribution 4.0 International or the GNU Lesser General Public License (LGPL) v. 2.0. We have also uploaded the GitHub data and code to Zenodo (DOI: 10.5281/zenodo.13845778).

## Ethics Statement

The animal experiments were performed with protocols approved by the University of Texas at Austin, Institutional Animal Care and Use Committee. The University of Texas at Austin animal management program is accredited by the Association for the Assessment and Accreditation of Laboratory Animal Care, International (AAALAC), and meets National Institutes of Health standards as set forth in the Guide for the Care and Use of Laboratory Animals (DHHS Publication No. (NIH) 85–23 Revised 1996).

## Acknowledgments

We are grateful to Jay Hinton for initially sharing the 10KSG dataset with us, to Ian Riddington for assistance with mass-spectrometry analysis of phospholipids, and Gerd Prehna and Larissa Diniz for discussions about enzymatic assay. Marcos Yoshinaga from PinguisLab for lipidomic data processing. Edgar Llontop and Diorge Paulo Souza for troubleshooting with protein purification. Cristiano G. Moreira for sharing strains. Alexandre Bruni for access to the microscope. Luize Nobrega Silva and Gustavo Chagas for technical support. This research was supported by Sao Paulo Research Foundation grant #2017-02178-2 to E.B-S, #2013/07937-8 to S.M, #2016/09047-8 and # 2021/10577-0 to R.F.S and #2021/10577-0 to T.W.C.S. Startup funds from MBS and CNS to E.B-S. This research was supported in part by an appointment to the National Library of Medicine Research Participation Program administered by the Oak Ridge Institute for Science and Education (ORISE) though an interagency agreement between the U.S. Department of Energy (DOE) and the National Library of Medicine. ORISE is managed by ORAU under DOE contract number DE-SC0014664. All opinions expressed in this paper are the author’s and do not necessarily reflect the policies and views of NIH, NLM, DOE, or ORAU/ORISE. This work is supported by the funds of the Intramural Research Program of National Library of Medicine at the National Institutes of Health, USA (to L.A. and G.G.N). This work utilized the NIH HPC Biowulf computer cluster.

## Supporting information

**S1 Fig. T6SS effector repertoire in the 10KSG dataset.** Each column indicates the presence or absence of a toxin as identified by the models developed in this study. Lines denote unique combinations of effectors. The histogram on the right shows the frequency of genomes in the 10KSG dataset containing each specific repertoire. The histogram at the bottom illustrates the frequency of genomes with at least one protein identified by the above HMM model. Effector activities are color-coded as described in Fig 2A.

S2 Fig. List of five most frequent T6SS effectors identified in each of the 149 *Salmonella* serovars contained in the 10KSG dataset. Colors indicate the effector activity as in Fig 2A.

**S3 Fig. Imm-Act1 is a two-transmembrane helices protein. (a)** Sequence alignment of Imm-Act1 homologs with orange rectangles indicating predicted transmembrane helices. Sequences are colored according to the Clustal X color scheme[97]. Consensus abbreviations: h, hydrophobic (A, C, F, I, L, M, V, W, Y); l, aliphatic (L, I, V); a, aromatic (F, W, Y); b, large residues (L, I, Y, E, R, F, Q, K, M, W); s, small residues (A, G, S, V, C, D, N); u, tiny residues (G, A, S); p, polar residues (S, T, E, D, K, R, N, Q, H, C); c, charged residues (D, E, H, K, R); +, positively charged residues (H, K, R); −, negatively charged residues (D, E). (B) Transmembrane helix prediction for Imm-Act1 using DeepTMHMM[98].

**S4 Fig. Recombinant protein purification used in enzymatic assay.** SDS–PAGE of recombinant proteins during purification steps to obtain purified protein for enzymatic assays. Affinity chomatography using Strep-Tactin sepharose to purify Tox-Act1 versions with C-terminus Strep-tag II: WT in (a) or C151A in (b). Recombinant proteins were purified from the soluble fraction. An additional step of washing with 1.5M urea was performed after the traditional washes to remove contamination with GroEL before elution with biotin. Additional bands were identified by mass-spectrometry to confirm identity. MK: marker; INS: insoluble; SOL: soluble; FT: flowthough. GroEL: chaperonin GroEL; Bccp: biotin carboxyl carrier protein; OmpF: outer membrane porin F.

**S1 Table. List of all T6SS toxin domains identified in this study in the 10K *Salmonella* genomes dataset.**

**S2 Table. List of all homologs collected by JackHMMER searches and used to build the phylogenetic tree shown in Fig 4A**.

**S3 Table. Genomic context of members of each permuted NlpC/P60 members shown in Fig 4A**.

**S4 Table. List of strains, plasmids and primers used in the study.**

**S5 Table. (a) MRM transition for LC-MS/MS method of lysophospholipids. (b) Internal standards used for lipidomics analysis in negative mode.**

**S1 Movie. Time-lapse microscopy of *E. coli* harboring pBRA SP-Tox-Act1 growing in media supplemented with 0.2% d-glucose.** Timestamp in hh:mm. Scale bar: 5 μm.

**S2 Movie. Time-lapse microscopy of *E. coli* harboring pBRA SP-Tox-Act1 growing in media supplemented with 0.2% l-arabinose.** Timestamp in hh:mm. Scale bar: 5 μm.

