## Supplementary figures and images for "Genome-directed study reveals the diversity of *Salmonella* T6SS effectors and identifies a novel family of lipid-targeting antibacterial toxins"

### Fig. S1

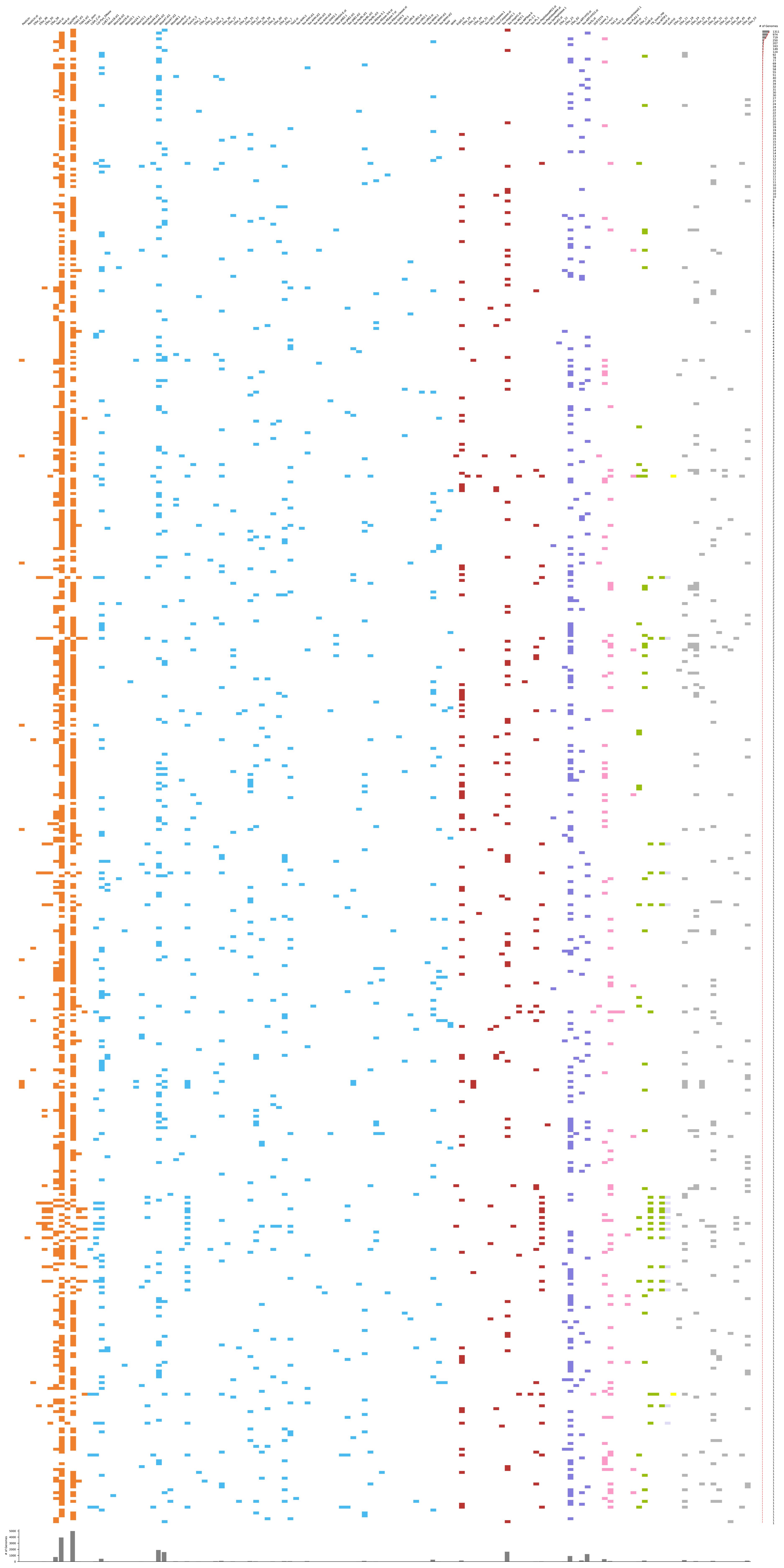

### Fig. S2

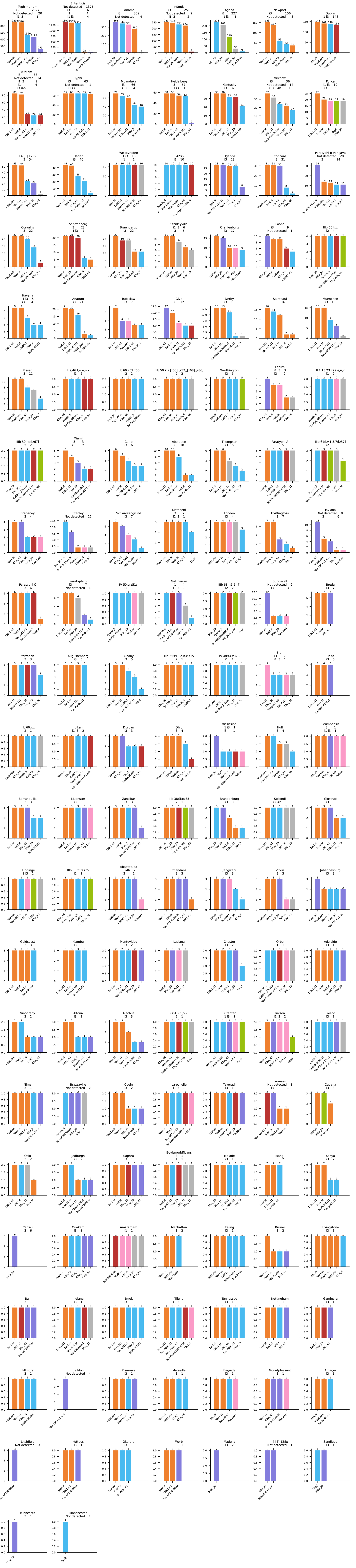

### Fig. S3

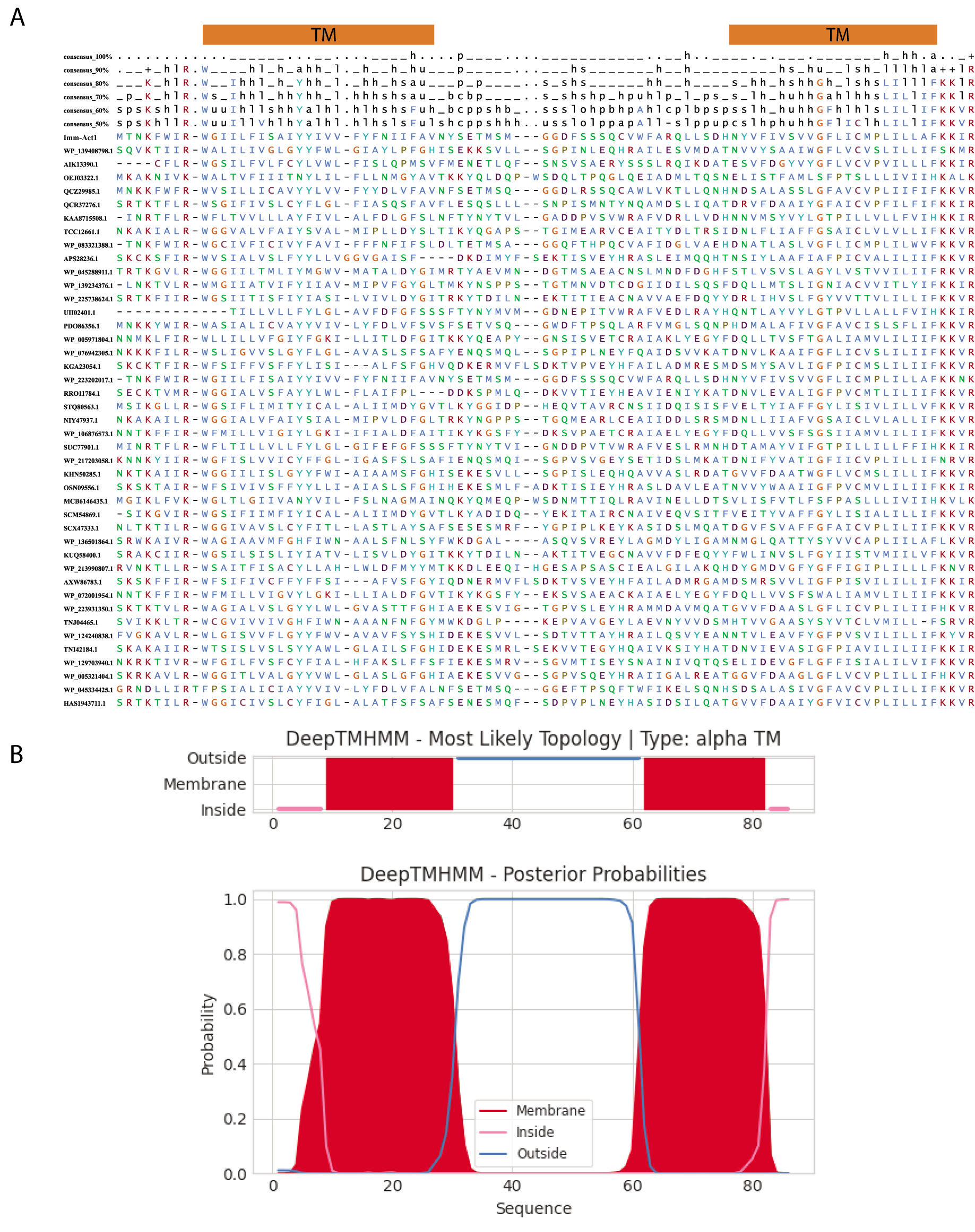

### Fig. S4

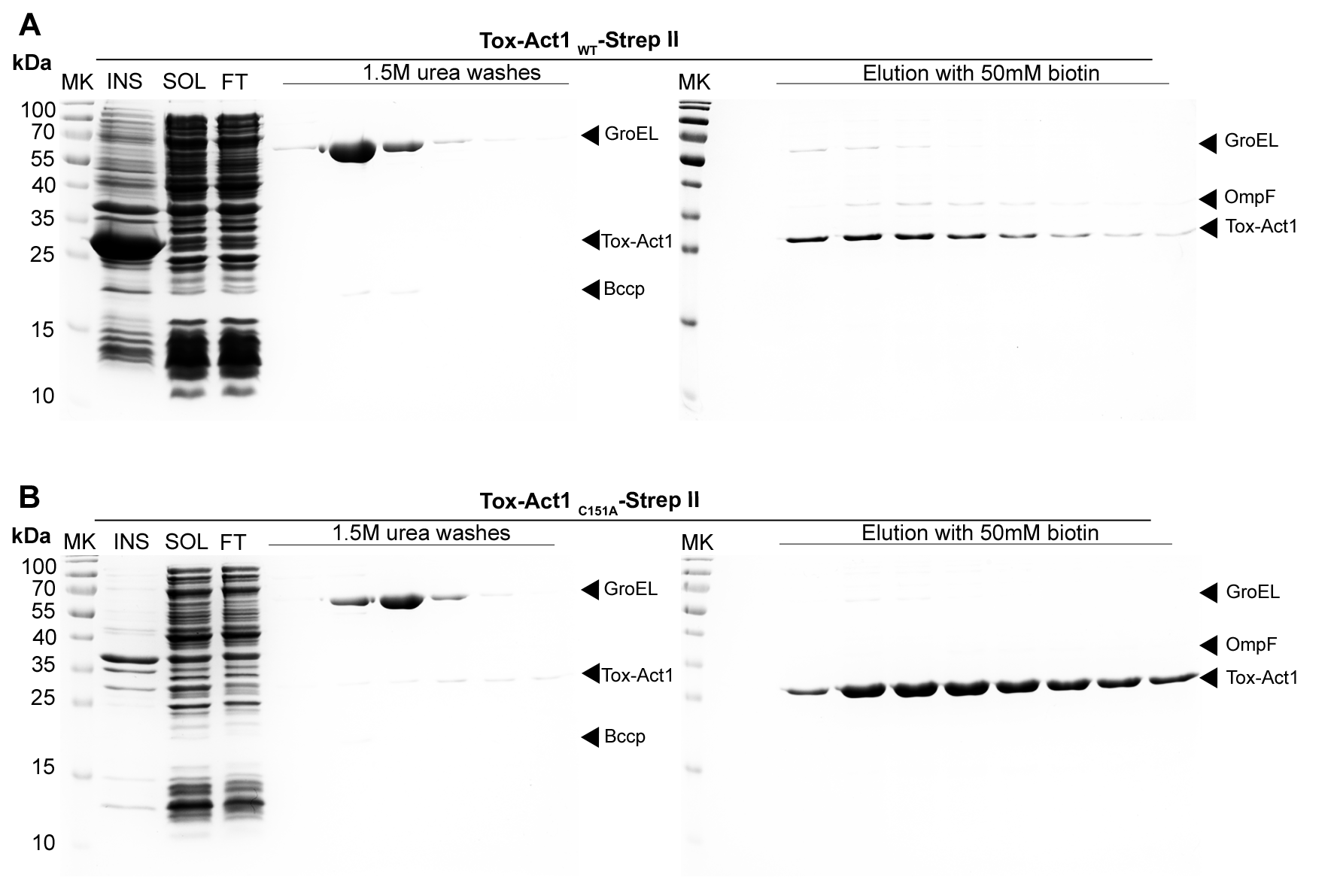
